# Arrestin-mediated Desensitization Enables Olfactory Discrimination in *C. elegans*

**DOI:** 10.1101/2021.05.02.439367

**Authors:** Daniel M. Merritt, Isabel MacKay-Clackett, Sylvia M. T. Almeida, Celina Tran, Safa Ansar, Derek van der Kooy

## Abstract

In the mammalian olfactory system, crosstalk among diverse olfactory signals is minimized through labelled line coding: individual neurons express one or few olfactory receptors among those encoded in the genome. Labelled line coding allows for separation of stimuli during mammalian olfactory signal transduction, however, in the nematode worm *Caenorhabditis elegans*, 1,300 olfactory receptors are primarily expressed in only 32 neurons, precluding this strategy. Here we report genetic, pharmacological and behavioural evidence that β-arrestin-mediated desensitization of olfactory receptors, working downstream of the kinase GRK-1, enables discrimination between intra-neuronal olfactory stimuli, but that this discrimination relies on quantitative, rather than qualitative differences in signalling. Our findings suggest that *C. elegans* exploits β-arrestin desensitization to maximize responsiveness to novel odors, allowing for behaviourally appropriate responses to olfactory stimuli despite the large number of olfactory receptors signalling in single cells. This represents a fundamentally different solution to the problem of olfactory discrimination than that which evolved in mammals, allowing for economical use of an extremely limited number of sensory neurons.

## Main Text

Olfaction is an enormously multidimensional sense^1^ through which signals generated by many odours are transduced through the nervous system to provide information about the external world. In mammals, crosstalk among these signals is minimized through the use of labelled line coding. Individual olfactory sensory neurons (OSNs) lining the olfactory mucosa stochastically express one or few olfactory receptors from the diverse repertoire encoded in the genome. Each OSN projects a single, unbranched axon, which passes through the cribriform plate to converge with the axons of other OSNs expressing the same receptor, forming one or few glomeruli specific to the receptor^2^. Within these glomeruli, OSN axons synapse onto the dendrites of mitral and tufted cells, which in turn project to the primary olfactory cortex. Thus, by ensuring physical isolation of nerves transducing signals from individual receptors, labelled line coding allows for clear separation of olfactory stimuli during early olfactory signal transduction.

The nematode worm *Caenorhabditis elegans* has a highly developed olfactory sense, with approximately 1,300 putative G protein-coupled receptor (GPCR) olfactory receptors^3^ encoded in its genome granting the ability to sense diverse chemical ligands^4^. Primary sensation of olfactory stimuli appears to be localized to approximately 32 sensory neurons^5^, with individual putative chemoreceptors usually being strongly expressed only in one bilateral pair of neurons^6,7^. Sensation of many odorants seems to depend primarily on the G protein α subunit ODR-3^4,8^, with the other G protein α subunits present in chemosensory neurons, GPA-2, GPA-3, GPA-5, GPA-6 and GPA-13, being able to partially compensate for its loss in experiments testing chemotaxis to some odorants^9^. The rich resource of olfactory receptors encoded in the worm’s genome presents a signalling conundrum: how can 1,300 receptors, all working through a limited set of partially redundant G proteins, maintain signalling identity (avoid crosstalk) when expressed in comparatively few chemosensory neurons^7,10^?

Evidence that crosstalk is avoided comes primarily from experiments in which an attractive odorant sensed exclusively by one neuron pair is placed on an agar plate containing a uniform concentration of a second attractive odorant, sensed exclusively by the same neuron pair. *C. elegans* is generally able to find the point source of the first odorant, despite the presence of the second, suggesting that olfactory discrimination between the two odorants is occurring. Importantly, when the uniform odour and the point odour are the same, animals are unable to locate the point, suggesting that attraction to the point does not result merely from a greater intensity of stimulus – instead, the response to the uniform odour has been “saturated”^4^.

We wondered if the apparent intra-neuronal olfactory discrimination exhibited by *C. elegans* in this paradigm might result from desensitization of the receptor or receptors required to sense the saturating odorant. Since the saturating odour is in the agar, and is in direct contact with the animal while the point odour can only reach it by diffusion, we reasoned that an activity-dependent process might affect receptors for the saturating odorant to a far greater extent than the point odorant, resulting in enhanced signalling through the latter relative to the former. This process would allow for chemotaxis toward the point by desensitizing the receptor or receptors responding to the saturating odorant, resulting in the receptors responsive to the point odorant being responsible for the majority of G protein signalling occurring in the neuron.

β-arrestins are the canonical desensitizers of activated GPCRs, and the β-arrestin family is represented in the *C. elegans* genome solely by the gene *arr-1*^11^. To determine whether arrestin-mediated desensitization was responsible for the apparent ability of worms to perform intra-neuronal olfactory discrimination, we first tested animals carrying *arr-1* null alleles in a saturation assay. *arr-1(ok401)* homozygous worms showed a severely abrogated ability to locate a point of isoamyl alcohol, sensed by the paired AWC neurons, within a saturating field of the AWC-sensed ^4^ odorant benzaldehyde. Conversely, in the absence of a saturating concentration of benzaldehyde, loss of arrestin resulted in only a very modest decrease in chemotaxis toward the point (Fig. 1a): this modest decrease may result from perturbation of chemosensory signalling, or may involve a role for ARR-1 in regulating non-chemosensory GPCRs required for optimal movement. These results were recapitulated in a second, probable null allele of *arr-1* (Fig. S1). Rescue of wild type *arr-1* under the *odr-3* promoter, driving expression primarily in AWC, mostly restored chemotaxis to a point of isoamyl alcohol within a saturating field of benzaldehyde (Fig. 1b), suggesting that expression of arrestin within AWC alone was sufficient to restore most discrimination between odours sensed by these neurons. We next sought to determine whether chemotaxis toward AWA-sensed odorants was similarly affected, and found that worms homozygous for *arr-1(ok401)* exhibited a significant, but not complete, block of chemotaxis to pyrazine within a saturating field of diacetyl, again with only a mild chemotactic deficit under unsaturated conditions (Fig. 1c). When tested in the presence of Barbadin, a small molecule inhibitor of the β-arrestin/AP2 complex, wild type worms displayed a similar inhibition of discrimination between AWC-sensed odours, and were unable to locate isoamyl alcohol in a saturating field of benzaldehyde (Fig. 1d), despite lacking any impairment of discrimination under unsaturated conditions (Fig. S2). Since Barbadin’s mechanism of action is specific to inhibition of the β-arrestin/AP2 complex, this suggests that arrestin enables olfactory discrimination by mediating endocytosis of the receptors for the saturating odour in clathrin-coated pits^12^.

**Fig. 1.**
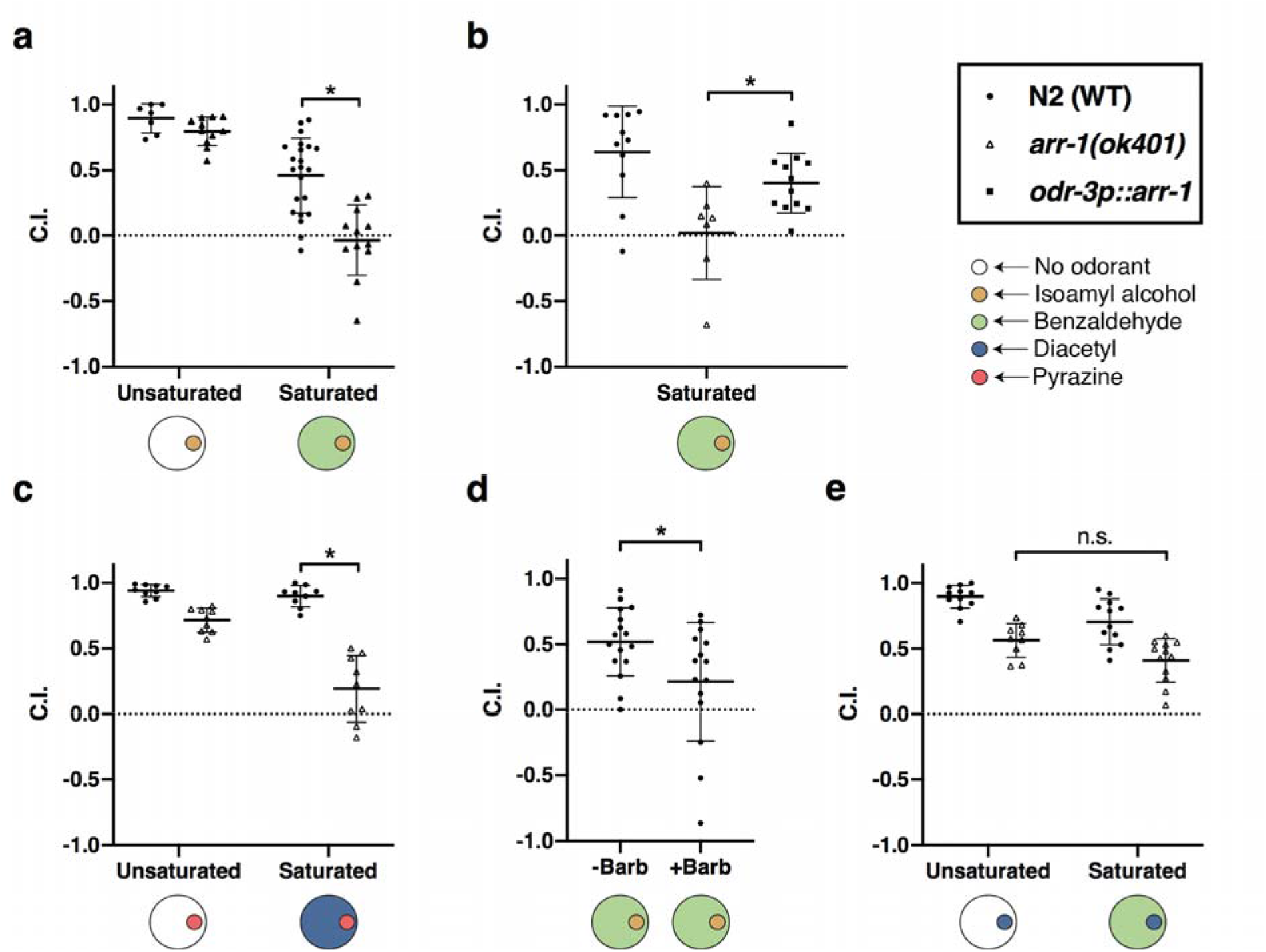
Olfactory discrimination depends on ARR-1. **a)** Chemotaxis of wild type N2 animals and *arr-1(ok401)* animals to a point of the AWC-sensed odorant isoamyl alcohol on unsaturated plates and plates containing a saturating concentration of the AWC-sensed odorant benzaldehyde. A two-way ANOVA revealed a significant effect of the interaction of strain and saturation condition (F=7.606, p<0.01), and a t-test revealed a significant difference between N2 and *arr-1(ok401)* in the benzaldehyde saturated conditions (t=4.997, p<0.01). **b)** Chemotaxis of wild type N2 animals, *arr-1(ok401)* animals and animals in which *arr-1* has been selectively rescued primarily in AWC to a point of isoamyl alcohol on plates containing a saturating concentration of benzaldehyde. A t-test revealed a significant difference between *arr-1(ok401)* and *odr-3p::arr-1* (t=2.56, p<0.05). **c)** Chemotaxis of wild type N2 animals and *arr-1(ok401)* animals to a point of the AWA-sensed odorant pyrazine on unsaturated plates and plates containing a saturating concentration of diacetyl. A two-way ANOVA revealed a significant effect of the interaction of strain and saturation condition (F=25.89, p<0.01), while a t-test revealed a significant difference between N2 and *arr-1(ok401)* in the diacetyl saturated conditions (t=7.96, p<0.01). **d)** Chemotaxis of wild type N2 animals to a point of the AWC-sensed odorant isoamyl alcohol on plates saturated with benzaldehyde, in the absence (-Barb) and presence (+Barb) of the β-arrestin inhibitor Barbadin. A t-test revealed a significant difference between the –Barb and +Barb groups (t=-2.27, p<0.05). **e)** Chemotaxis of wild type N2 animals and *arr-1(ok401)* animals to a point of the AWA-sensed odorant diacetyl on unsaturated plates and plates containing a saturating concentration of the AWC-sensed odorant benzaldehyde. A two-way ANOVA revealed no significant interaction between strain and saturation condition (F=0.171, p>0.05).

Our hypothesized mechanism of intra-neuronal olfactory discrimination should, if correct, play a minimal role in inter-neuronal olfactory discrimination, where labeled line coding is possible because of physical separation of signalling, allowing for the avoidance of crosstalk. To test this prediction, we evaluated *arr-1(ok401)* animals for chemotaxis to a point of an AWA-sensed odorant, diacetyl, in a saturating concentration of the AWC-sensed odorant benzaldehyde. As expected, no saturation-dependent chemotactic deficit was apparent in *arr-1(ok401)* animals (Fig. 1e), suggesting that arrestin-mediated desensitization does not play a role in discrimination in contexts in which labelled line coding is sufficient to distinguish between odorants.

Previous reports have determined saturating concentrations in this paradigm by finding the minimum concentration of the odorant that prevented *C. elegans* from locating a point of the same odorant^4^. We wondered if arrestin might, by desensitizing activated receptors, create a ceiling on receptor signalling potential, resulting in saturation at lower levels. To test this, we evaluated *arr-1(ok401)* animals for chemotaxis to a point of benzaldehyde in a saturating concentration of benzaldehyde. Consistent with previous reports, we found that wild type N2 animals were unable to find the point of benzaldehyde, however loss of arrestin resulted in a modest but significant increase in the ability to find the point (Fig. 2a). This result, demonstrating an increase in the dynamic range of benzaldehyde-induced signalling, is a novel prediction of our hypothesis, further supporting a role for arrestin in constraining the dynamic range of response to ubiquitously present odours.

**Fig. 2.**
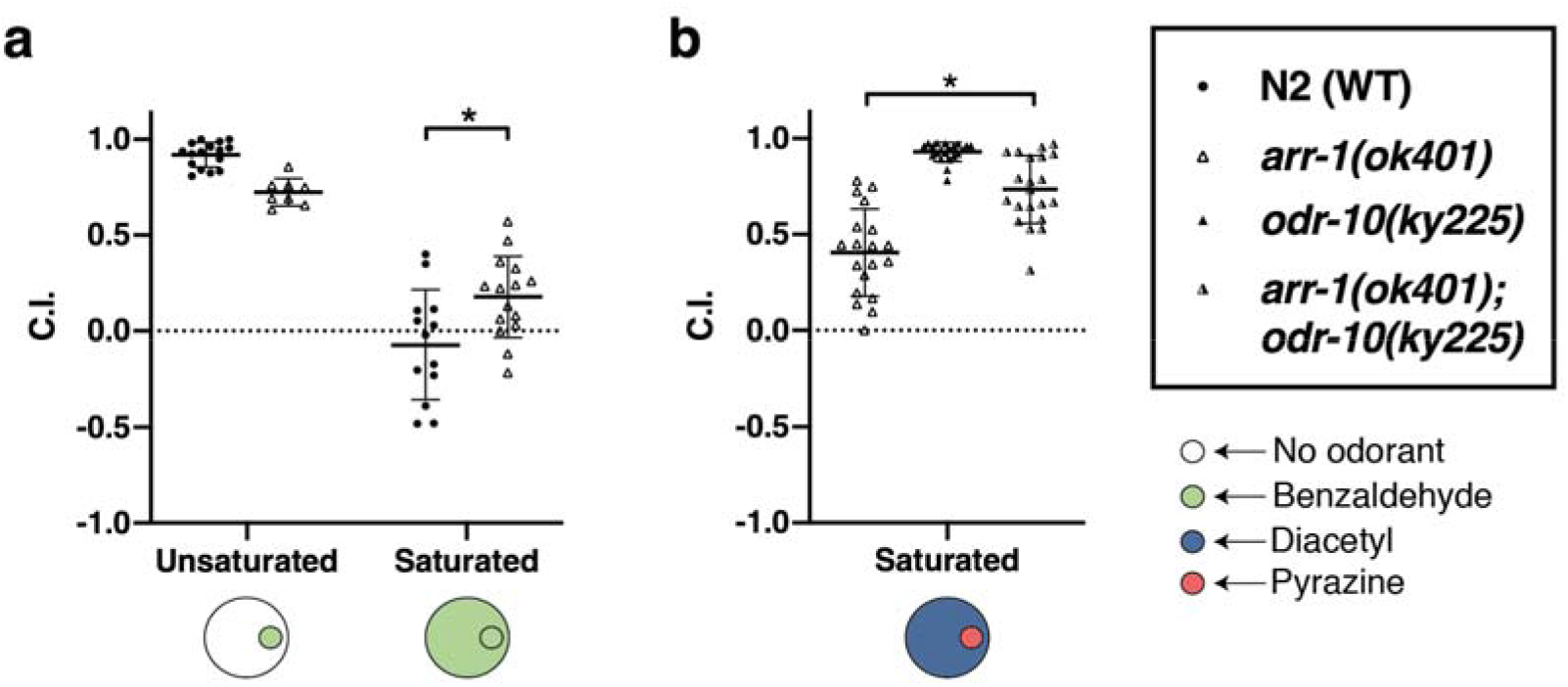
Experimental support for model predictions. **a)** Chemotaxis of wild type N2 animals and *arr-1(ok401)* animals to a point of benzaldehyde on both unsaturated agar plates, and plates containing a saturating concentration of benzaldehyde. A two-way ANOVA revealed a significant interaction between strain and saturation condition (F=16.60, p<0.01), and a t-test indicated a significant difference between N2 and *arr-1(ok401)* in the benzaldehyde saturated condition (t=-2.58, p<0.05). **b)** Chemotaxis of *arr-1(ok401), odr-10(ky225)*, and *arr-1(ok401); odr-10(ky225)* animals to a point of pyrazine on plates containing a saturating concentration of diacetyl. A t-test revealed a significant difference between *arr-1(ok401)* and *arr-1(ok401);odr-10(ky225)* animals (t=-5.07, p<0.01).

If discrimination of point odorants from saturating odorants occurs via arrestin-mediated desensitization of the receptors corresponding to the saturating odorants, then genetic elimination of these receptors should suppress the discrimination defects we observed in *arr-1* mutant animals. We exploited a mutant in *odr*-*10*, the sole *C. elegans* receptor responsive to low concentrations of diacetyl^13^, to test our proposed mechanism of intra-neuronal discrimination by examining chemotaxis to a point of pyrazine in a saturating concentration of diacetyl. *odr-10(ky225)* animals, which carry a deletion in the majority of the *odr-10* gene, have wild type responses to pyrazine but no chemotactic responses to low to moderate concentrations of diacetyl^13^. As expected, *odr-10(ky225)* and *arr-1(ok401);odr-10(ky225)* animals showed minimal approach to diacetyl (Fig. S3). In subsequent experiments testing approach to pyrazine in a saturating concentration of diacetyl, *odr-10(ky225)* animals exhibited robust chemotaxis to the point, while *arr-1(ok401)* animals showed only weak approach. In *arr-1(ok401);odr-10(ky225)* double mutants, however, approach to pyrazine was largely restored, indicating suppression of *arr-1* by *odr-10* (Fig. 2b). This finding – that elimination of the ability to sense one odorant can enhance the ability of the animal to sense a second – is predicted by our model, but would be surprising in a labelled line model, and thus further supports our hypothesis.

In mammals, arrestin-mediated desensitization occurs following phosphorylation of the activated receptor by G protein-coupled Receptor Kinases (GRKs)^14^. Although the *C. elegans* genome contains two predicted GRKs, *grk-1* and *grk-2*, loss of function mutations in these genes have been found to result in surprising phenotypes. While in mammals loss of *GRK3* (expressed in the olfactory epithelium) results in olfactory desensitization deficits similar to those caused by loss of function mutations in arrestin^15,16^, loss of *C. elegans grk-2* appears to completely abrogate chemosensation^17^, while the only phenotype known to result from mutations in g*rk-1* is an alteration in dopamine-mediated swimming-induced paralysis^18,19^. We wondered whether arrestin-mediated olfactory discrimination might constitute a sensitized assay capable of revealing subtle chemosensory deficits in *grk-1* mutant animals. When tested for chemotaxis to a point of isoamyl alcohol in a saturating field of benzaldehyde, *grk-1* mutants showed severe chemotactic deficits. Unlike in *arr-1* mutants, no chemotaxis deficit to isoamyl alcohol in unsaturated conditions was observed, suggesting that loss of *grk-1* may result in either more specific (effecting fewer GPCRs), or less complete (effecting the same GPCRs to a lesser extent), inhibition of desensitization than loss of *arr-1* (Fig. 3).

**Fig. 3.**
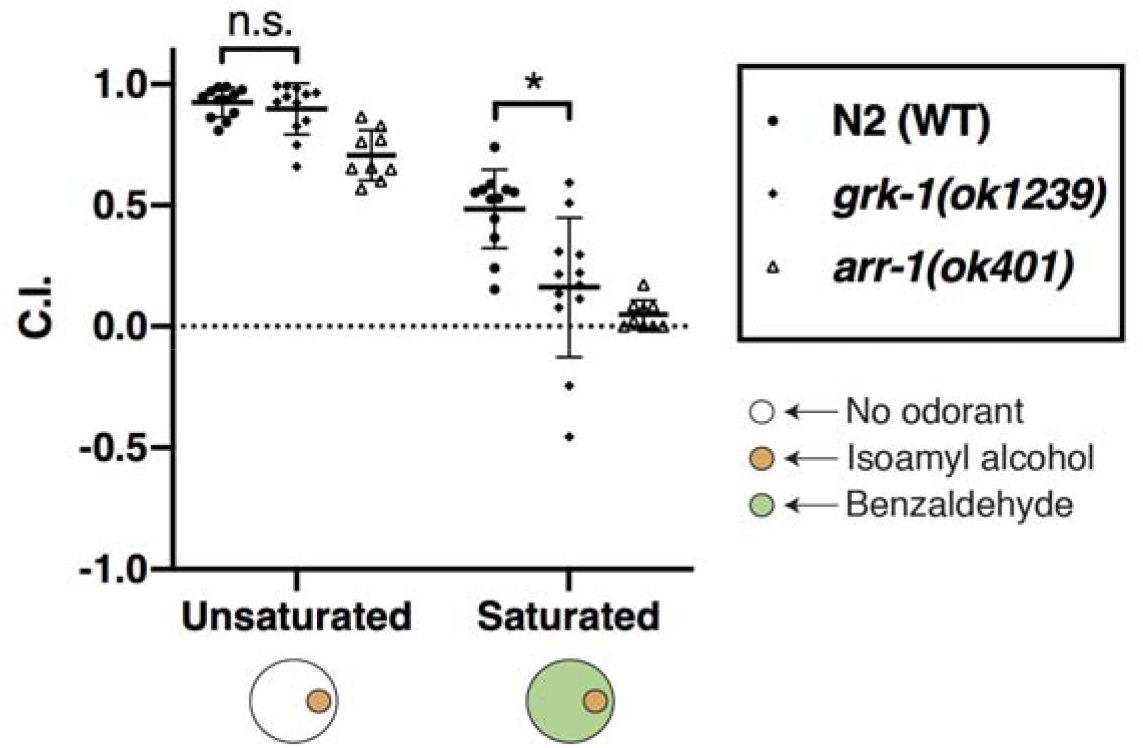
GRK-1 activity is required for olfactory discrimination. Chemotaxis of wild type N2 animals, *arr-1(ok401)* and *grk-1(ok1239)* animals to a point of isoamyl alcohol on plates containing a saturating concentration of benzaldehyde. A two-way ANOVA revealed a significant interaction between strain and saturation condition (F=5.695, p<0.01), and a t-test indicated a significant difference between N2 and *grk-1(ok1239)* in the benzaldehyde saturated condition (t=-3.40, p<0.01), but no significant difference between N2 and *grk-1(ok1239)* in the unsaturated condition (t=-0.76, p > 0.05).

Nematodes and mammals face a similar problem in ensuring signal segregation between the many chemosensory receptors providing their sense of olfaction. While mammals have solved this problem by expressing olfactory receptor genes in a one or few genes per neuron fashion, our findings suggest that nematodes have enacted a radically different solution, in which many chemosensory receptors are expressed within a given neuron, but discrimination is enabled through desensitization of ubiquitously active receptors (Fig. 4). In mammals, labelled line coding preserves a qualitative difference between olfactory stimuli deep into the nervous system, however in nematodes an initial qualitative difference at the receptors is converted by arrestin to a quantitative difference in signalling, which, by resulting in selective responses to odorants, nonetheless results in qualitatively different behavioural responses. By allowing for expression of far more olfactory receptors than the worm has neurons to express them in, this strategy greatly enhances the olfactory repertoire of *C. elegans*.

**Fig. 4.**
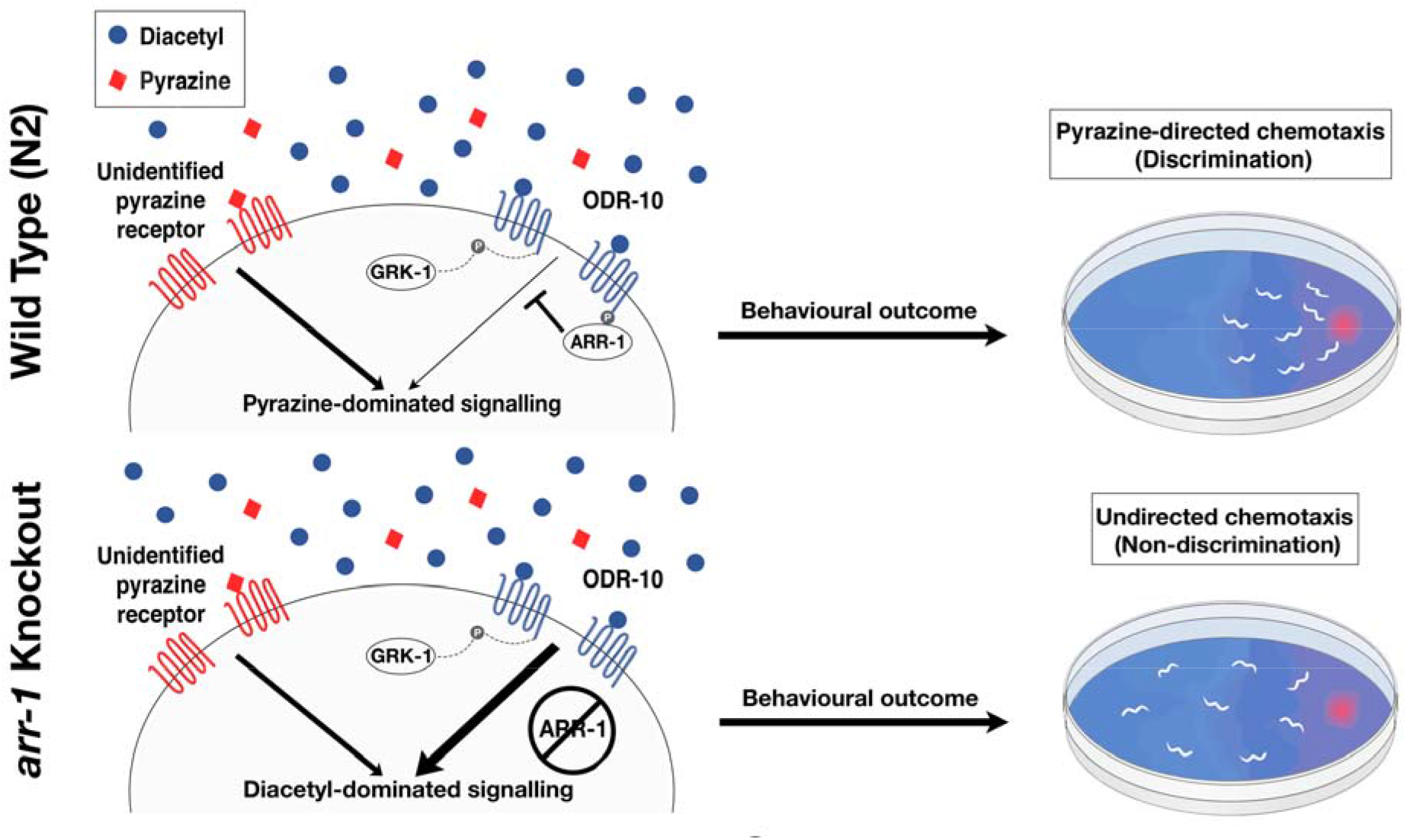
Diagram of discrimination model. In olfactory neurons of wild type N2 animals (top), ARR-1 acts to desensitize the receptors for saturating odorants, here shown as the AWA-sensed odorant diacetyl, leaving only signalling from the point odorant, here shown as the AWA-sensed odorant pyrazine, to determine chemotactic behaviour. In *arr-1* mutant animals (bottom), ubiquitous signalling from the receptor for the saturating odorant overwhelms signalling from the point odorant, preventing chemotaxis toward it.

## Acknowledgments

Some strains were provided by the CGC, which is funded by NIH Office of Research Infrastructure Programs (P40 OD010440). The authors thank Srinidhi Krishnakumar for technical assistance.

## Funding

This research was supported by NSERC RGPIN Grant 8319, and by NSERC CREATE in Manufacturing, Materials, Mimetics.

## Author contributions

Behavioural experiments were performed by D.M.M., I.M.C., C.T. and S.A. Genetic manipulations were performed by S.M.T.A. The manuscript was written by D.M.M. and I.M.C., and all authors reviewed it. D.v.d.K. supervised the work and edited the manuscript.

## Competing interests

Authors declare no competing interests.

## Data and materials availability

All data is available in the main text or the supplementary materials.

## Notes

### Competing Interest Statement

The authors have declared no competing interest.

