## Supplemental Material for "Arrestin-mediated Desensitization Enables Olfactory Discrimination in *C. elegans*"

**Materials and Methods:**

*C. elegans* Strains and Maintenance

Animals were maintained at 20°C on NGM plates and fed *Escherichia coli* OP50. All experiments were performed at 20°C.

The *C. elegans* strains N2, RB660 *arr-1(ok401)*, RB1194 *grk-1(ok1239)* and CX3410 *odr-10(ky225)* were provided by the CGC. UT1313 *arr-1(ok401);odr-10(ky225)* was produced by crossing RB660 and CX3410. *arr-1(vs96)* and the *Ex[odr-3p::arr-1] arr-1(ok401);lin-15(n765ts)* rescue line were kindly provided by Jeffrey Benovic.

Preparation of Worms for Behavioural Assays

Worms were age-synchronized using a sodium hypochlorite bleaching protocol. Briefly, eggs were isolated following treatment of worms with bleach solution and incubated on a rocker overnight at 20°C in ~2.5 mL M9 buffer to allow hatching, after which approximately 1000 L1 larvae were plated on an NGM plate seeded with *E. coli* OP50 and allowed to grow for 53.5-54 hours at 20°C (to the young adult stage).

Young adult worms were first washed from their cultivation plates with ~1.5 mL M9 buffer and collected into 1.5 mL microcentrifuge tubes with a Pasteur pipette. Worms were washed twice with additional volumes of M9 to remove any remaining bacteria. During the wash steps, the largest worms were selected: worms were re-suspended in buffer and only those worms that settled within ~1 min. were collected. Worms were starved in 1.5 mL of M9 buffer for ~90 minutes before the assay.

Behavioural Assays

Population chemotaxis assays were performed as previously described (*4*). Briefly, assay plates were 10 cm Petri plates containing 10 mL of assay agar (1.6% agar, 5 mM $KH_2PO_4$ pH 6.0, 1 mM $CaCl_2$, 1 mM $MgSO_4$). All attractant points were diluted in anhydrous ethanol to the following concentrations: benzaldehyde (Sigma-Aldrich) 1:200, isoamyl alcohol (Bioshop) 1:10, pyrazine (Sigma-Aldrich) 1:100, diacetyl (Fluka) 1:1000.

Assays testing detection of one attractant in the saturating presence of another were performed on plates to which undiluted attractant was added in a 1:10000 ratio directly to the agar before pouring. Agar was first liquefied in a microwave and then cooled to ~60°C by incubating in a water bath for at least 1 hour. Both odorant-saturated plates and unsaturated control plates were sealed with Parafilm after pouring.

Approximately 50-200 worms were transferred to the centre of an assay plate using a calibrated glass micropipette in ~10 µL of M9 buffer. 1 µL of 1 M sodium azide (Sigma-Aldrich) was spotted at opposite sides of the plate, 1 cm from the edge, to paralyze worms at the spots where attractant odorant would be spotted. Worms were allowed to settle in the small volume of M9 buffer at the centre of the assay plate for 2 minutes before 1 µL attractant odorant diluted in ethanol and 1 µL of 100% ethanol were spotted on opposing sides of the plate, on top of the sodium azide points. A corner of a Kimwipe twisted into a point was then used to dry the worms. Worms were observed under the microscope during drying to ensure the surface of the agar was not broken with the Kimwipe; plates with any large breaks introduced at the origin during drying were not scored. Plates were again sealed with Parafilm and left undisturbed for 60 minutes, after which a chemotaxis index (C.I.) was calculated as (number of worms within 1 cm of the ethanol counter attractant subtracted from the number of worms within 1 cm of attractant) / (total number of worms on the plate). Any worms having not moved from a defined 1 cm by 0.5 cm rectangle where they were initially placed were considered injured and omitted from this calculation. Any plates with fewer than 20 total worms were excluded from analysis.

Barbadin Experiments

Worms were added to the centre of the test plate in an approximately 10 uL drop of 1 mM Barbadin (Toronto Research Chemicals) (or DMSO in control conditions) dissolved in 2% pluronic acid (F-127, Sigma Aldrich) in M9. Barbadin was prepared as a stock solution at 100 mM in DMSO. In addition to Barbadin in the drop at the centre of the plate, the agar in these experiments contained 100 µM Barbadin (or DMSO in control conditions) and 2% pluronic acid.

Statistical Analysis

No statistical test was performed to predetermine sample sizes. Mean C.I. and the SEM were calculated using R. In experiments with multiple independent variables, comparisons between C.I. were made by two-way ANOVA and follow-up t-tests to determine between-strain differences within a testing condition. In experiments with a single independent variable, comparisons were made by t-test alone. In experiments with more than one test, t-test results were adjusted using Bonferroni correction. Differences were considered significant when $p<0.05$.

**Supplemental Figures:**

**Fig. S1.**

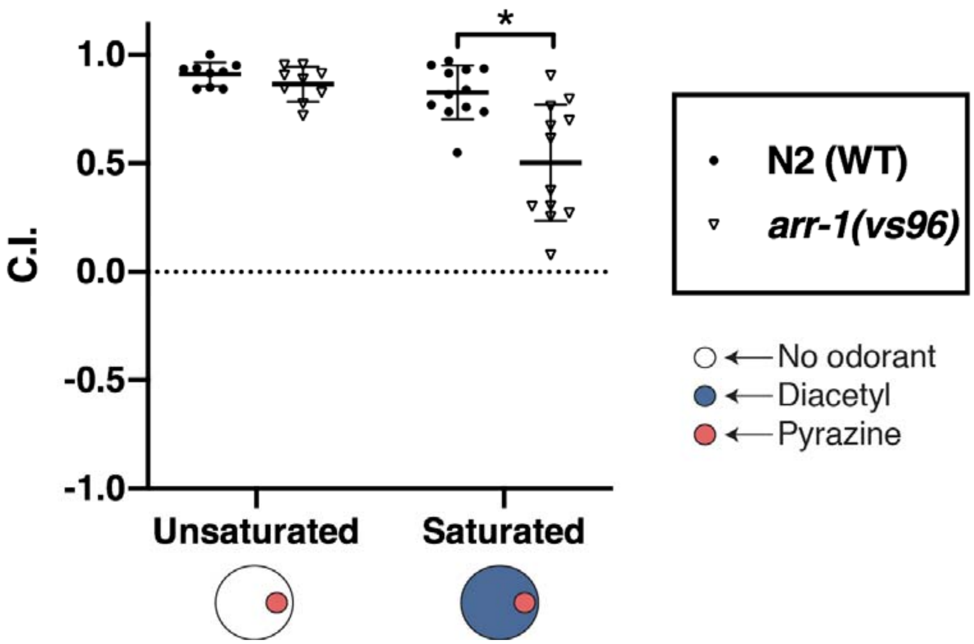


**Fig. S1 *arr-1(vs96)* exhibits an AWA olfactory discrimination deficit similar to *arr-1(ok401)*.** Chemotaxis of wild type N2 animals and *arr-1(vs96)* animals to a point of the AWA-sensed odorant pyrazine on unsaturated plates and plates containing a saturating concentration of the AWA-sensed odorant diacetyl. A two-way ANOVA revealed a significant interaction between strain and saturation condition ($F=7.294$, $p<0.05$), and a t-test indicated a significant difference between N2 and *arr-1(vs96)* in the diacetyl saturated condition ($t=3.80$, $p<0.01$).

**Fig. S2**

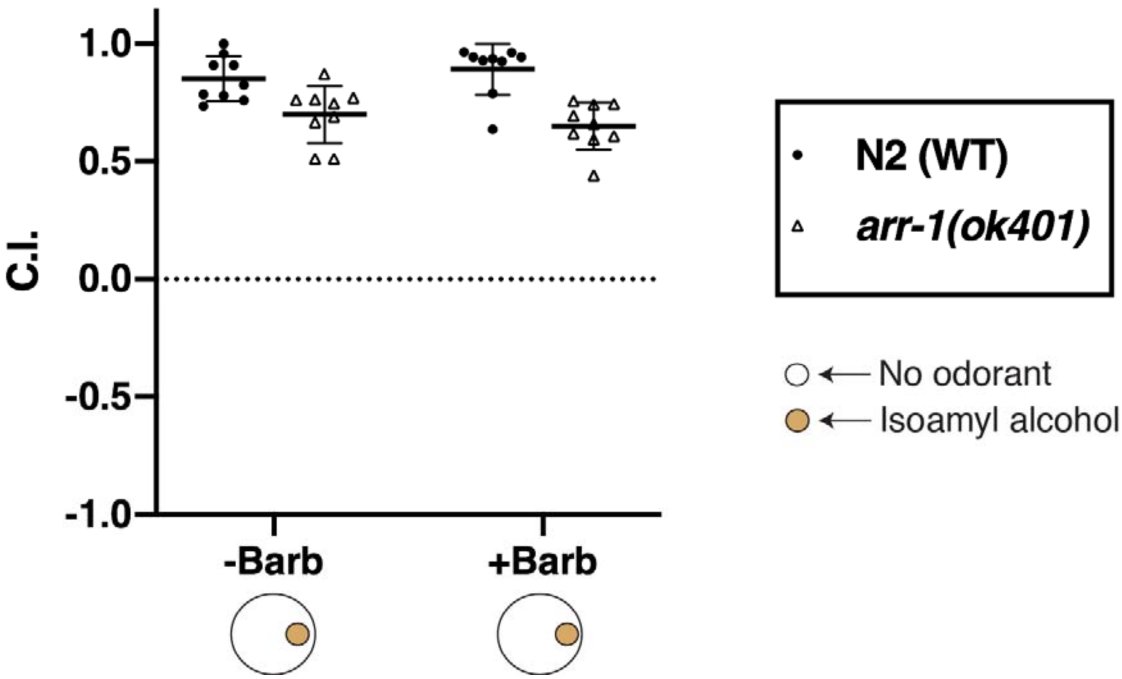


**Fig. S2 Barbadin has no effect on unsaturated chemotaxis.** Chemotaxis of N2 and *arr-1(ok401)* animals to a point of isoamyl alcohol on plates with and without Barbadin. A two-way ANOVA revealed no effect of drug presence ($F=0.012$, $p > 0.05$).

**Fig. S3.**

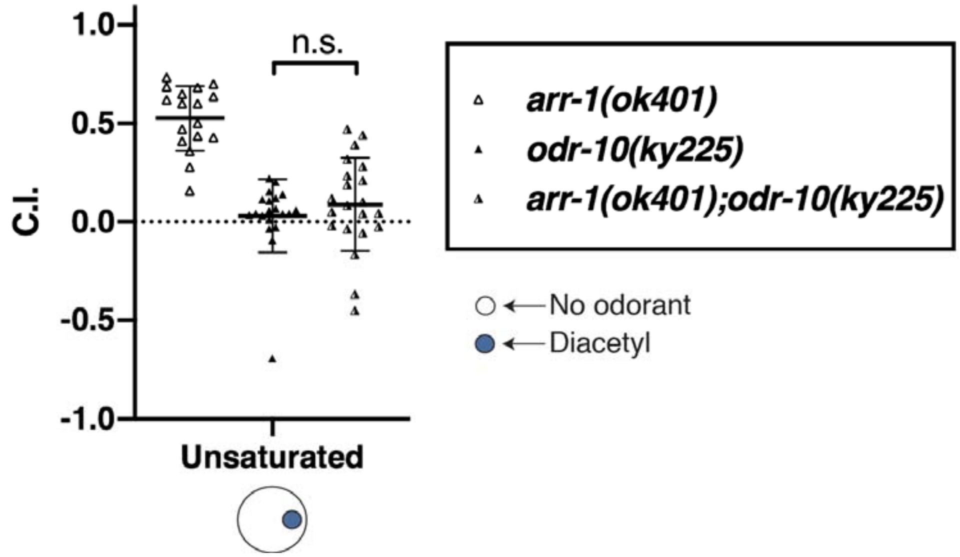


**Fig. S3 *arr-1(ok401);odr-10(ky225)* double mutant animals and *odr-10(ky225)* single mutant animals display similar diacetyl chemotaxis.** Chemotaxis of *arr-1(ok401)*, *odr-10(ky225)*, and *arr-1(ok401); odr-10(ky225)* animals to a point of diacetyl on unsaturated plates. A t-test between *odr-10(ky225)* and an *arr-1(ok401);odr-10(ky225)* double mutant revealed no significant difference ($t=-0.87$, $p>0.05$).
